# Computations performed by shadow enhancers and enhancer duplications vary across the Drosophila embryo

**DOI:** 10.1101/396457

**Authors:** Clarissa Scholes, Kelly M. Biette, Timothy T. Harden, Angela H. DePace

## Abstract

Transcription of developmental genes is controlled by multiple enhancers. Frequently, more than one enhancer can activate transcription from the same promoter in the same cells. In these cases, how is regulatory information from multiple enhancers combined to determine the overall expression output of their shared promoter? To investigate this question, we quantified nascent transcription driven by a pair shadow enhancers, each individual of the pair, and their duplications in *Drosophila* embryos using live imaging. This set of constructs allows us to quantify the “computation” made by the pairs of enhancers: their combined output expression as a function of the expression that they drive separately. We show that the computation performed by these shadow enhancers and duplications varies across the expression pattern, implying that how their activities are combined depends on the transcriptional regulators bound to the enhancers in different parts of the embryo. Characterizing the computation made by multiple enhancers is a critical first step in developing conceptual and computational models of gene expression at the locus level, where multiple enhancers collaborate.

## INTRODUCTION

Developmental genes are controlled by many enhancers, some of which can drive overlapping patterns of gene expression in space and time (Dukler et al., 2016; Dunipace et al., 2011; Frankel et al., 2010; Ghiasvand et al., 2011; Hay et al., 2016; Hoch et al., 1991; Hong et al., 2008; Jeong et al., 2006; Lam et al., 2015; McBride et al., 2011; Osterwalder et al., 2018; Perry et al., 2011; Prazak et al., 2010; Zeitlinger et al., 2007). If enhancers are active in different cells and work independently of one another, their overall transcriptional output should be predicted by superimposing their activities. But “shadow” enhancers, which drive spatiotemporally overlapping expression of the same gene (Hong et al., 2008), produce patterns and levels of expression that cannot be predicted from their separate activites (Bothma et al., 2015; Dunipace et al., 2011; Prazak et al., 2010). When two or more enhancers are simultaneously active, we do not yet understand how their activities are integrated to determine the level and timing of gene expression.

Shadow enhancers are pervasive at developmental genes in vertebrates and invertebrates (Barolo, 2011; Cannavò et al., 2016; Kvon et al., 2014), where they are thought to improve the precision and robustness of expression (Barolo, 2011; Frankel, 2012; Frankel et al., 2010; Lam et al., 2015; Perry et al., 2010). While they drive similar expression patterns, shadow enhancers are not functionally identical and can collaborate to fine-tune the spatial or temporal boundaries of gene expression (El-Sherif and Levine, 2016; Perry et al., 2011; Prazak et al., 2010). Their partially-redundant behaviour may also enable shadow enhancers to buffer against environmental or genetic perturbations; deletion of a shadow enhancer may have no effect under normal developmental conditions and only be revealed under conditions of stress (Frankel et al., 2010; Perry et al., 2010). The mechanisms by which shadow enhancers might enable robust gene expression remain unclear. One suggestion is that having more than one enhancer increases the likelihood of gene activation at any given time (Perry et al., 2010); another is that shadow enhancers help to maintain the level of gene expression above a critical threshold (Ghiasvand et al., 2011; Lam et al., 2015). Deciphering the role of shadow enhancers in development will require elucidating the underlying molecular mechanisms that control how they collaborate.

While we know a great deal about the molecular mechanisms of transcription initiation and elongation (Fuda et al., 2009), we know much less about how enhancers regulate these processes, let alone how multiple enhancers control them through interactions with the same promoter. Enhancers contact the promoter to activate transcription, often from a distance of many kilobases (Spitz, 2016), and recent live imaging experiments showed that transcription of a gene in fact only occured when the promoter and its distant enhancer were in close physical proximity (Chen et al., 2018). Enhancers also make contact with one another, as demonstrated by chromatin capture experiments (Ghavi-Helm et al., 2014), and this type of assay has also shown that more than one enhancer is able to physically contact the promoter at a time on a single DNA template (Oudelaar et al., 2018). Reciprocally, live imaging has revealed that a single enhancer can activate two promoters at a time (Fukaya et al., 2016). Together, these studies paint a picture of “many-by-many” interactions between enhancers and promoters.

One route to unraveling the molecular mechanisms of how multiple enhancers interact with a single promoter is to measure their “computation”: their combined output expressed as a function of that driven by each enhancer acting alone. For instance, a pair of enhancers may together drive expression that is equal to, greater-than or less-than the sum of their separate activities (Bothma et al., 2015). This computation reflects the underlying molecular mechanisms, and therefore quantifying it is a critical first step in developing conceptual and computational models of expression at the locus level, where multiple enhancers are together responsible for dictating gene expression.

On the basis of live imaging data, shadow enhancers have been proposed to activate the promoter one at a time, and this was formalized in a version of the two-state model for transcription extended to two enhancers (Bothma et al., 2015). In this model, if enhancers only contact the promoter infrequently then they are unlikely to get in one another’s way together will drive additive expression. On the other hand, when enhancers frequently activate the promoter they will compete for access and drive transcription at a less-than-additive rate. Bothma et al. showed that where enhancers drive low levels of expression, their combined effects tend to be additive –– as would be predicted if low expression were the result of infrequent contact with the promoter. Meanwhile highly-expressing enhancers together produce less-than-additive expression, consistent with their contacting the promoter frequently and thus competing for access to it. In this model, frequency of contact with the promoter determines competition between enhancers. However, since it’s not yet possible to measure contact directly in gene loci with closely-spaced enhancers, the strength of expression from each enhancer was used as a proxy for their contact frequency.

To investigate how information from multiple active enhancers is combined, we quantified the computation performed by a pair of shadow enhancers driving expression of *Krüppel* in *Drosophila melanogaster* blastoderm-stage embryos. Previous studies have measured expression from a shadow enhancer pair and compared it to that driven by each enhancer separately using *in situ* hybridization in fixed embryos (Dunipace et al., 2011; Perry et al., 2011; Prazak et al., 2010; Wunderlich et al., 2015) and, more recently, using the MS2 system in live embryos (Bothma et al., 2015; El-Sherif and Levine, 2016). Live imaging is particularly powerful for studying the computations made by enhancers and their cognate promoters, as early *Drosophila* development is rapid and expression changes quickly in space and time. Our study is distinguished from previous efforts in three important ways. First, we measure expression driven by the individual *Krüppel* enhancers, duplicates of each, and the shadow enhancer pair. The duplicated enhancer constructs enable us to distinguish the effects of simply having two enhancers that activate the same promoter from effects of the shadow enhancer pair. Second, because expression can vary in subtle ways depending on the distance of an enhancer relative to the promoter, all of our constructs are distance controlled. Third, we analyze the computation made by the various enhancer arrangements, and compare it to the predictions of the model described above, in which enhancers compete to activate the promoter (Bothma et al., 2015).

*Krüppel* is well-suited to test the competition model because its central domain of expression is driven by two shadow enhancers (Figure 1A,B), one of which drives low levels of expression and the other high (Figure 1E). Duplicating lowly-expressing and highly-expressing enhancers is the minimal perturbation with which to test whether expression level, as a proxy for enhancer-promoter contact frequency, is what primarily determines the computation made by simultaneously-active enhancers. Comparing the behaviour of duplicated enhancers to that of the shadow enhancer pair also enables us to determine whether binding the same or different sets of transcription factors affects how the enhancer’s activities are integrated. Since distance between enhancer and promoter influences expression dynamics (Fukaya et al., 2016), we compare expression driven by *Krüppel’s* pair of shadow enhancers, and duplications of each, to expression driven by the appropriate distance-controlled single enhancer constructs (Figure 1C).

**Figure 1:**
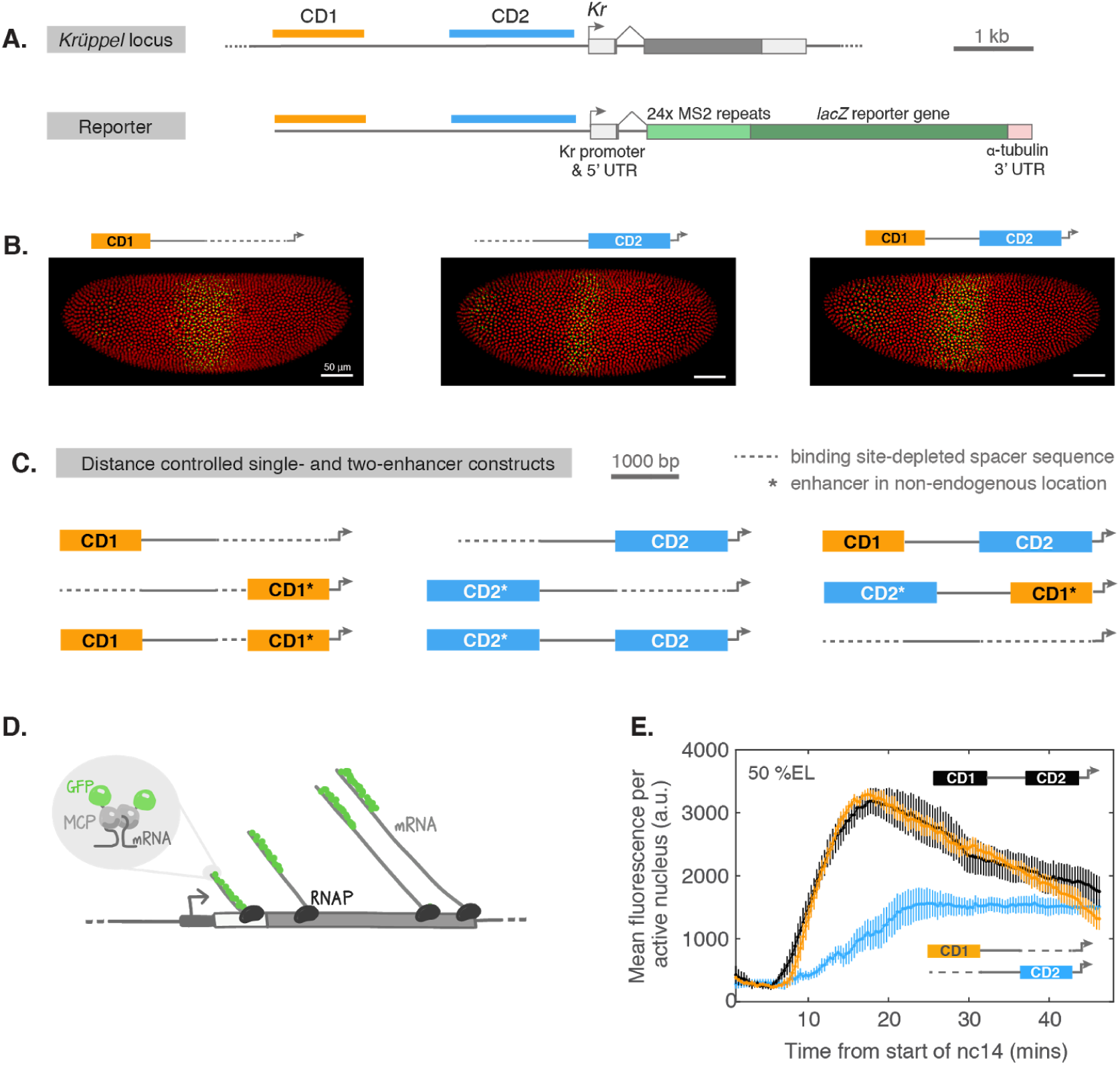
Constructs for live imaging of nascent transcription. **(A).** *Krüppel* (*Kr*) locus showing the two blastoderm enhancers, CD1 (orange) and CD2 (blue). The *Krüppel* reporter used here drives expression of an MS2 cassette followed by a *lacZ* reporter gene and α-tubulin 3’UTR, under the control of the endogenous *Krüppel* promoter and 5’UTR. **(B).** The blastoderm enhancers of *Krüppel* drive overlapping expression patterns in the central region of the embryo during nuclear cycle 14 (nc14). Representative maximum projection images showing nascent transcription (MCP-GFP; green) and nuclei (histone-RFP; red) in mid-nc14 (anterior left, dorsal up). **(C).** Reporter constructs used to investigate the behavior of enhancer duplications. * indicates an enhancer is at its non-endogenous location relative to the promoter. Dotted lines represent synthetic “neutral” sequences, computationally designed to be depleted for binding sites of transcription factors active in patterning the blastoderm embryo. These sequences were used to maintain the endogenous spacing of the enhancers and promoter in the reporter constructs. **(D).** Live imaging of nascent transcription using MS2 system. A sequence encoding 24x MS2 repeats was incorporated into the 5’ of the reporter gene. When it is transcribed, the sequence forms RNA stem loops that are bound by a constitutively expressed MCP-GFP fusion protein. As a train of polymerases transcribe through the reporter, GFP builds up at the locus and nascent transcripts become visible. Once a transcript dissociates from the gene and diffuses away it is no longer discernible. **(E).** *Krüppel* shadow enhancers drive differing levels and dynamics of expression over time in nuclear cycle 14 (CD1, orange; CD2, blue; CD1-CD2, black). Plot shows mean fluorescence per active nucleus in a single anterior-posterior bin of 2.5% embryo length (EL) covering between 50-52.5 % EL (see also Figure S1). Error bars are standard error of the mean across multiple embryos. Each time point is separated by 20 seconds. CD1, n = 5 embryos; CD2, n = 6; CD1-CD2, n = 6.

Our results demonstrate that the computation performed by two enhancers is not simply a constant function of their individual expression level. At the edges of the pattern, the computation is different in the anterior than it is in the posterior. By switching the order of the shadow enhancers we also show that all relevant information is not contained within the enhancers; rather, additional information is contained in their relative positions to one another in the locus, which influences the computation they perform. Our results reinforce that molecular mechanisms to control gene expression must be considered at multiple levels, from individual transcription factors, to collections of transcription factors in enhancers, to collections of enhancers in complex loci. In particular, the mechanisms operating at the locus level remain to be clearly elucidated.

## RESULTS

To measure the dynamic computations performed by combinations of two enhancers across the *Drosophila* embryo, we generated a comprehensive set of constructs containing one or two copies of the blastoderm enhancers of *Krüppel* (Figure 1A). The constructs drive expression of an MS2 reporter cassette that enabled us to image nascent transcription in developing embryos (Garcia et al., 2013; Lucas et al., 2013) (see Methods for details). *Krüppel’s* enhancers, CD1 and CD2 (Hoch et al., 1990), lie within 4 kb upstream of the promoter and drive spatio-temporally overlapping patterns of expression in the developing embryo (El-Sherif and Levine, 2016; Hoch et al., 1990; Perry et al., 2011) (Figure 1B). Since the distance between enhancer and promoter affects expression levels (Fukaya et al., 2016), we made control constructs to measure expression from each enhancer in single copy at the alternate location (i.e. CD1 at the proximal position, and CD2 at the distal position, denoted CD1* and CD2*, respectively; Figure 1C). We followed the sequence definitions for the *Krüppel* enhancers established by Perry et al (2011)(Table S1), which were based on the binding of *Krüppel* regulators measured in ChIP-chip assays (Li et al., 2008). To maintain the endogenous distances between regulatory elements, and to ensure that we did not accidentally introduce binding sites for regulators of *Krüppel*, we replaced each enhancer in the single-enhancer constructs with an equivalent length of synthetic neutral sequence depleted of binding sites for transcription factors involved in patterning the blastoderm embryo (Estrada et al., 2016) (Methods; Table S2).

### Single copies of enhancers drive expression influenced by position relative to promoter

In order to determine the computation carried out by combinations of two enhancers –– i.e. the expression driven by two enhancers together as a function of their separate activities –– we must first accurately measure their separate activities. Our single-enhancer constructs control for quantitative variations in expression at different positions of the enhancer relative to the promoter, allowing us to discern the computation made by different enhancer combinations.

*Krüppel* expression driven by the shadow enhancers pair (the CD1-CD2 construct) rises rapidly during nc14 to a peak at ∼18 minutes from the start of the cycle, before declining again (Figure 1E). The level of expression is set primarily by CD1, while CD2 turns on slightly later and more gradually to a lower peak level that, unlike CD1 expression, is maintained through the remainder of nc14 (Figure 1E). CD2 also drives a narrower pattern than CD1 (Figure 2G) and represses the activity of the distal CD1 enhancer in the posterior of the pattern (El-Sherif and Levine, 2016). We report the mean expression per actively-transcribing nucleus (Figure 2A, E), and the fraction of nuclei that are transcribing (Figure 2B, F); together these dictate the overall mean expression level driven by each enhancer. We note that the relative level of expression driven by CD1 differs between our results and a previous study, which may be due to differences in the sequence definitions of the enhancers and/or promoters between our two studies (see Discussion).

**Figure 2:**
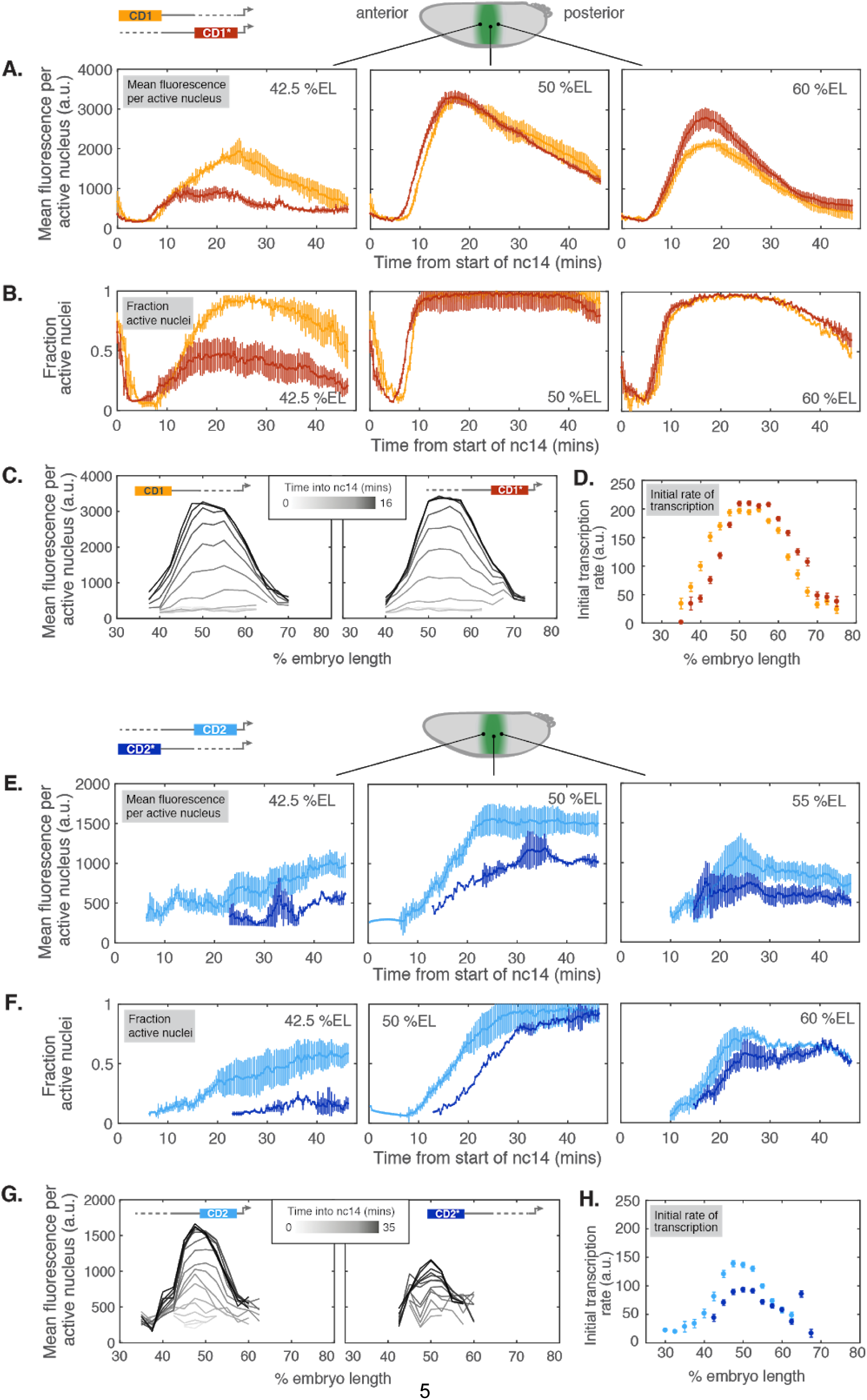
Single copies of enhancers drive expression influenced by position relative to promoter. **(A).** Comparing mean expression over time in nuclear cycle 14 (nc14) driven by the CD1 enhancer at its endogenous position (CD1, orange) and at the promoter (CD1*, red). Shown is mean fluorescence per active nucleus (a.u.) within bins in the anterior, middle and posterior of the expression pattern. Each bin is 2.5% embryo length (EL) in width. Error bars show the standard error of the mean and data points are 20 secs apart. CD1, n = 3 embryos; CD1* n = 6. **(B).** Fraction of total nuclei in the indicated anterior-posterior bin that are actively driving transcription, over time in nc14. Nuclei are counted as active if there is a detectable spot of MS2 fluorescence. Total number of nuclei in a given bin in nc14 is ∼30. Error bars show the standard error of the mean across multiple embryos. **(C).** Evolution of the expression pattern driven by CD1 (left) and CD1* (right) in the first 16 mins of nc14. Line traces show the mean fluorescence in active nuclei across the pattern and show time points ∼100 secs apart. **(D).** Initial transcription rate across the patterns driven by CD1 and CD1*, estimated by finding the maximum derivative of the initial rise in fluorescence for each transcriptionally active nucleus; data points show mean across all active nuclei across a number of embryos for each anterior-posterior bin, +/-standard error. **(E).** Mean fluorescence per active nucleus (a.u.) driven by the CD2 enhancer at its endogenous position (blue) and upstream of the promoter (CD2*, indigo) across nc14. CD2, n = 6 embryos; CD2* n = 4. **(F).** Fraction of active nuclei driven by CD2 and CD2* over time in nc14. **(G).** Evolution of the expression pattern driven by CD2 (left) and CD2* (right) over the first 35 mins of nc14. The window of time shown is longer than in the equivalent plots in (C) because the CD2 enhancer takes longer to start driving expression and to reach its peak level. **(H).** Initial transcription rate across the expression patterns driven by CD2 and CD2* (see D).

The expression driven by both *Krüppel* shadow enhancers is affected by their distance from the promoter. Moving CD1 next to the promoter from its endogenous distance ∼2.8 kb upstream, reduces expression in the anterior of the *Krüppel* pattern while increasing it slightly in the posterior (Figure 2A,C,D). When CD2 is moved away from its endogenous position adjacent to the promoter, expression turns on later and to a lower level across the entire pattern (Figure 2E,F,H).

### *Krüppel* enhancer duplications and shadow enhancer pair drive different computations over the anterior-posterior axis

We next examined how the expression driven by each pair compares to the sum of expression driven by the appropriate single distance-controlled enhancers (Figure 3). We used additive expression as the baseline for comparison since this is the predicted outcome if two enhancers independently activate transcription without competing with one another for the promoter (Bothma et al., 2015). Each combination of enhancers drives sub-additive expression, consistent with this model (Figure 3A-C).

**Figure 3:**
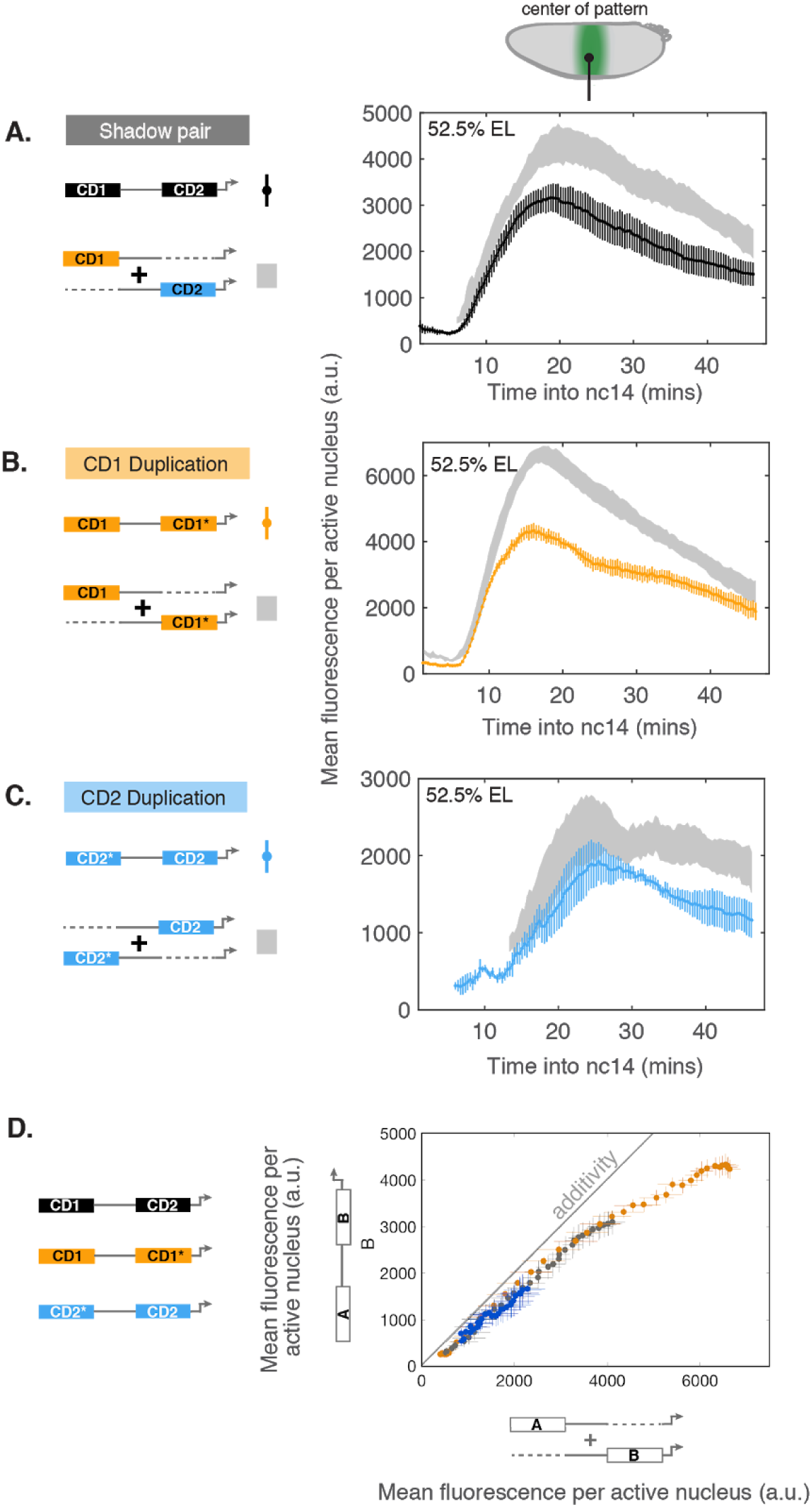
Duplications and shadow enhancers of Krüppel drive largely sub-additive expression. **(A).** Mean expression over time in nuclear cycle 14 (nc14) driven by the shadow enhancer pair (black), compared to the sum of expression driven by each distance-controlled individual enhancer (grey). Mean fluorescence per active nucleus +/- standard error at approximately the center of the expression pattern, in the bin at 52.5% embryo length (EL). CD1, n = 3 embryos; CD2, n = 6; CD1-CD2, n = 6. **(B).** Comparison of expression from the duplicated CD1 construct (orange) compared to the sum of expression from CD1 and CD1* alone (grey). CD1, n = 3 embryos; CD1*, n = 6; CD1-CD1*, n = 3. **(C).** Comparison of expression from the duplicated CD2 construct (blue) to the projected sum of expression driven by CD2* and CD2 (grey). CD2, n = 6 embryos; CD2*, n = 4; CD2*-CD2, n = 5. **(D).** In the center of the expression pattern, the two-enhancer constructs (shadow enhancer pair, CD1 duplication and CD2 duplication) follow the same relationship to additivity over a wide range of expression levels. This holds over ∼10% of embryo length in the anterior-middle of the pattern (not shown). Plot shows comparison of expression driven by each two-enhancer construct to the sum of expression driven by the constituent enhancers (‘A’ and ‘B’). Time points shown are from the period during which mean expression is increasing to a peak: 3-18 mins from the start of nc14 (shadow enhancer pair and CD1 duplication) and 10-25 mins (CD2 duplication). Shown is the mean fluorescence per active nucleus, +/- standard error, from the middle of the expression patterns (bin at 52.5% embryo length). The standard error on the x-axis is the summed error in the fluorescence driven by each constituent enhancer. Grey line indicates additivity.

In order to directly compare the computations performed by the shadow enhancers and enhancer duplications, we plotted the expression level from each pair against the sum of the expression driven by the relevant distance-controlled single enhancers (Figure 3D). In the center of the *Krüppel* pattern, these pairs of enhancers follow a strikingly similar trend with respect to additivity over a wide range of expression levels (Figure 3D). Consistent with the predictions of the competition model, when the enhancers drive low expression levels the output expression is close to additivity (exemplified by the duplication of CD2), while when they drive high levels the output is sub-additive (exemplified by the CD1 duplication).

However, at the borders of the expression pattern, the computation differs between the anterior and posterior of the pattern and between pairs of enhancers (Figure 4). For example, CD1 and CD2 together drive expression that is close to additive in the anterior of the pattern (grey, Figure 4A), but much less than additive in the posterior (black, Figure 4A). This behaviour reflects a documented interaction between the two enhancers, in which CD2 represses the activity of CD1 in the posterior part of the pattern (El-Sherif and Levine, 2016).

**Figure 4:**
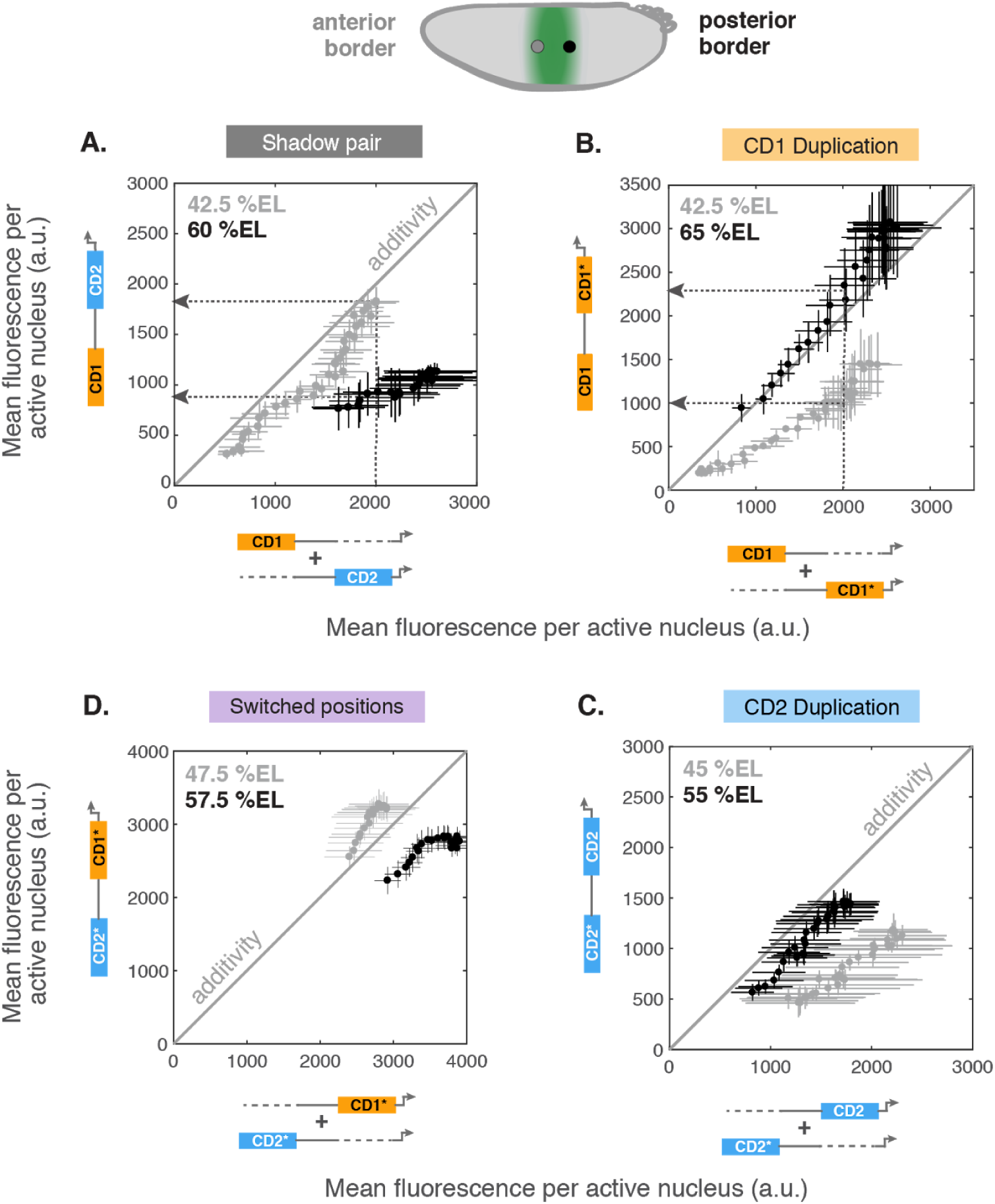
Computations performed by two enhancers vary over the anterior-posterior axis. We compare mean fluorescence per actively transcribing nucleus driven by each combination of two enhancers to the sum of the constituent enhancers’ expression at the anterior (grey) and posterior (black) borders of the pattern. Each datapoint is a different time point (at 20 sec intervals) over the first 20 minutes of nuclear cycle 14. Note that we only plot data for time points at which all three relevant constructs (the single enhancers and both enhancers together) drive expression. **(A).** The shadow enhancer pair drives approximately additive expression in the anterior (at 42.5% embryo length, grey), but sub-additive expression in the posterior of the pattern (at 60% embryo length, black). The dotted arrows highlight how the same summed expression level from the two enhancers alone (on the x-axis, where CD1+CD2 = 2000 a.u.) yields different expression level outputs at different anterior-posterior positions in the embryo. **(B).** Expression from the CD1 duplication is approximately additive in the posterior of the pattern (black) and sub-additive in the anterior (grey; note that this is the opposite trend to that displayed by the shadow pair). For a given summed level of expression from CD1 and CD1* on the x-axis, the duplication yields different expression levels at different anterior-posterior positions (dotted arrows). **(C).** Expression from the CD2 duplication compared to the sum of expression driven by CD2 and CD2* separately. Data shown in (C) is from the first 25 mins of nc14. **(D).** When the positions of the shadow enhancers are switched (CD2*-CD1* construct), expression is additive (or even super-additive) in the anterior (grey), while it is less than additive in the posterior of the pattern (black), following the same trend as when the two enhancers are in their endogenous arrangement.

The difference in the computation performed across the *Krüppel* pattern also exists when the two enhancers involved are identical (Figure 4B,C). Consider the CD1 duplication; at a given level of ‘input’ expression (x-axis), it is possible to get different output expression levels (y-axis). This trend is construct-specific: the CD1 duplication is closer to additive in the posterior, whereas the shadow enhancer pair is closer to additive in the anterior (compare grey to grey in Figures 4A and B). Finally, the difference in the computation carried out at the anterior and posterior borders of the pattern persist when the positions of the shadow enhancers are switched in the reporter construct (compare Figure 4D to 4A). This variation in computation at the boundaries result from different transcription factors controlling expression in the anterior versus posterior of the pattern (see Discussion).

### Repression of CD1 by CD2 does not require CD2 to be adjacent to the promoter

The CD2 enhancer refines the *Krüppel* expression pattern by inhibiting activity of the CD1 enhancer at the posterior boundary (Figure 5A; El-Sherif et al. 2016). We wondered whether CD2’s repressive activity relies on either its promoter-proximal location or its positioning between CD1 and the promoter. To test this, we switched the positions of the two enhancers and found that it does not: the CD2 enhancer is as effective at preventing activation by CD1 in the posterior of the *Krüppel* expression pattern from 2.4 kb upstream of the promoter and on the opposite side of CD1 from it (Figure 5B-C).

**Figure 5:**
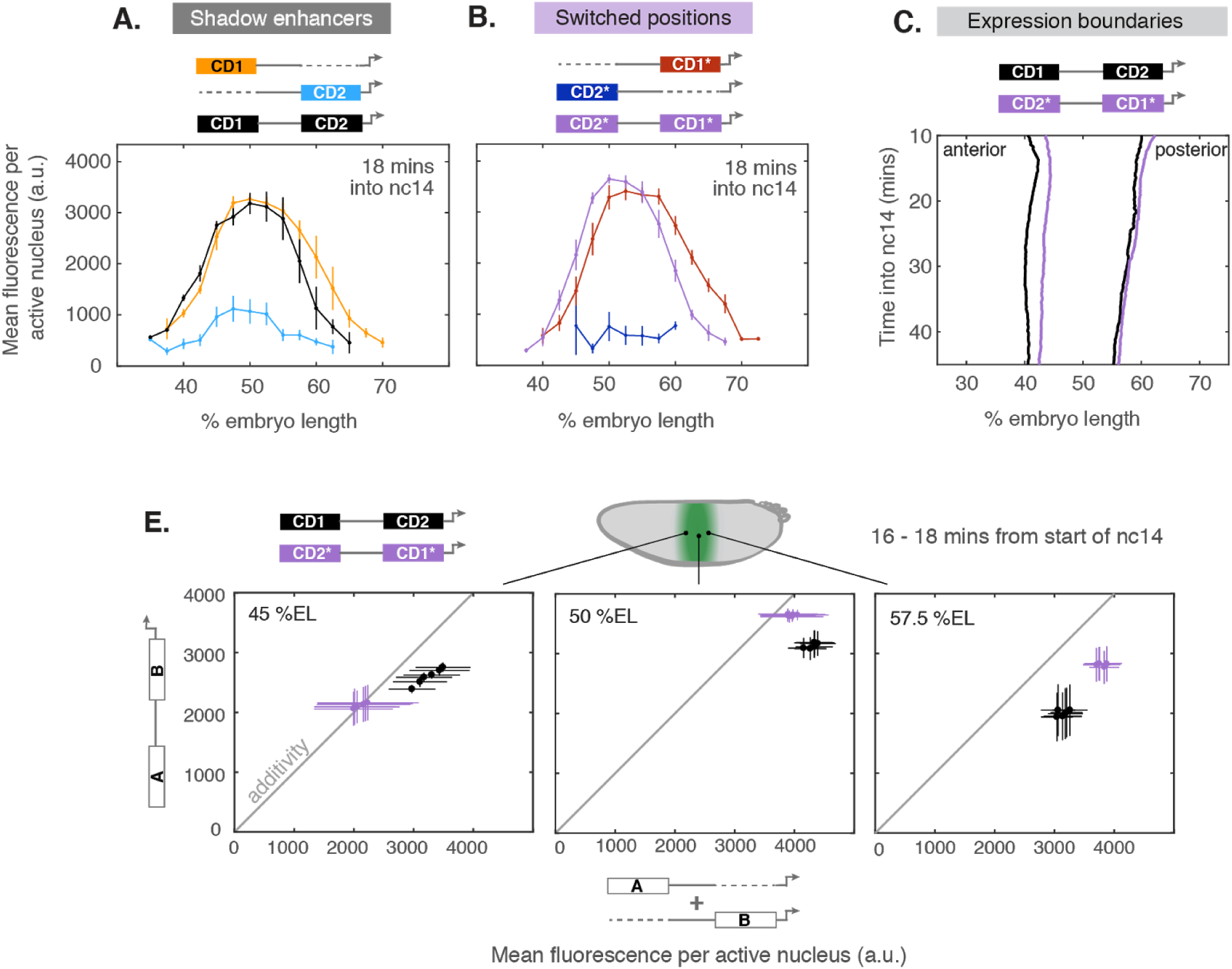
Regulatory sequence arrangement influences computation performed by two enhancers. **(A).** When both shadow enhancers drive expression together (CD1-CD2, black), the CD2 enhancer (blue), represses the activity of CD1 (orange) in the posterior of the *Krüppel* pattern. Plotted is mean expression per active nucleus across the *Krüppel* expression pattern at a single time point, 18 minutes into nuclear cycle 14 (nc14). Error bars are standard error of the mean across multiple embryos. **(B).** When the positions of the shadow enhancers are switched (CD2*-CD1*, purple), the CD2* enhancer (indigo) still represses the activity of CD1* (red) effectively in the posterior of the pattern. **(C).** Comparison of the anterior and posterior boundary positions driven by the shadow enhancer pair in their endogenous (black) and switched (purple) locations over time in nc14. The posterior boundary remains the same irrespective of the order of the enhancers. However, the anterior border driven by construct containing the enhancers in their switched positions (CD2*-CD1*) is shifted backwards by ∼2.5% embryo length relative to that driven by the endogenous construct (CD1-CD2). Anterior and posterior boundaries were defined by fitting a Gaussian to the expression pattern at each time point and finding the positions at half maximum expression (see Figure S1). **(D).** Comparison of the computation performed by the shadow enhancers in their endogenous (black) and switched positions (purple) across the *Krüppel* pattern. Mean fluorescence from each two-enhancer construct (y-axis) is plotted against the sum of the mean expression from the appropriate single-enhancer constructs (x-axis). Data are shown from six time points between 16-18 minutes from the start of nc14. Error bars are standard error of the mean across multiple embryos.

### Regulatory sequence arrangement influences computation by two enhancers

Finally, we asked whether the computation performed by the shadow enhancer pair depends only on the sequences of the enhancers, or whether the position of the enhancers relative to one another also influences how they interact. We switched the locations of the pair of shadow enhancers, and compared the computions between the endogenous and switched arrangements. This analysis was possible because we measured the output of individual enhancers controlled for their distance from the promoter. These single enhancer controls enable us to separate the effect of individual enhancer location on expression from the effect of enhancer arrangement on the computation itself. The regulatory sequences in the endogenous and switched constructs are essentially identical (Figure 1C; see Methods). If the computation they carry out is solely a function of the sequence of each enhancer, then we would expect them to combine in the same way regardless of their arrangement. However, switching their positions impacts the computation, as was evident when we compared CD1-CD2 and CD2*-CD1* constructs (Figure 5E). These data indicate that the organisation of the gene locus influences not only the level and timing of expression driven by each enhancer alone (Figure 2A,E), but also how the two enhancers work together to determine the overall level of expression.

## DISCUSSION

We quantified nascent transcription in developing *Drosophila* embryos to study how a promoter combines information from multiple concurrently-active enhancers. Duplicating strongly-expressing and weakly-expressing shadow enhancers of the *Krüppel* gene enabled us to test the idea that enhancers compete for access to the promoter. It also allowed us to investigate whether there are differences in how inputs from multiple enhancers are “computed” depending on whether they contain the same or different combinations of transcription factor binding sites. Our data are consistent with *Krüppel*’s enhancers competing to activate the promoter one at a time, but they also indicate that the complement of transcription factors bound to the enhancers in different parts of the embryo, and the arrangement of enhancers relative to one another, influence the computation the enhancers perform.

### Sequence context matters in quantitative measurement of gene expression

#### Distance between enhancer and promoter affects expression

We found that the distance between enhancer and promoter quantitatively affects expression level and timing (Figure 2), contrary to the canonical definition of enhancers as position- and orientation-independent (Banerji et al., 1981). This effect differs between the two enhancers, and it is non-uniform across the anterior-posterior axis of the embryo. For instance, CD1 drives lower expression in the anterior of the pattern when it is positioned next to the promoter. This implies that *Krüppel* is more sensitive to the repressors that position its anterior boundary, Giant and Hunchback (Jaeger, 2010), when those repressors are bound close to the promoter. The broad decrease in expression driven by CD2 when it is moved away from the promoter is consistent with previous work that found expression to decline with distance between enhancer and promoter (Fukaya et al., 2016).

It is common for studies to use reporter constructs in which enhancers are placed immediately adjacent to the promoter, or to delete enhancers from bacterial artificial chromosomes without replacing the deleted sequence, which alters the spacing between regulatory elements. Our results demonstrate the importance of controlling for the endogenous spacing of enhancers and promoters when making quantitative expression measurements. When introducing spacers, we used computationally designed synthetic sequences that lack predicted binding sites (Estrada et al., 2016), but it remains possible that even this “neutral sequence” has an effect on expression. In future work, it would be useful to test many different spacers to control for this possibility.

#### Differences in relative expression level compared to previous studies may stem from differences in enhancer and promoter definitions

In our constructs, CD1 and CD2 drive different relative levels of expression compared to constructs reported by two previous studies, including one from our own lab (El-Sherif and Levine, 2016; Wunderlich et al., 2015). Both Wunderlich et al. and El-Sherif & Levine reported lower peak expression from CD1 alone, a level similar to that driven by their CD2-only constructs.

The constructs used in both previous studies differed from those in the current work. Wunderlich et al. (2015) used the minimal *even-skipped* promoter and did not control for distance between CD1 and the promoter. Meanwhile, El-Sherif & Levine (2016) used different sequence definitions of the enhancers, a minimal core *Krüppel* promoter; and the CD1-only construct was not controlled for the distance of this enhancer from the promoter. When we compared the predicted binding sites across all of our constructs and those of El-Sherif & Levine, we saw that their CD1-only construct alone does not contain a consensus motif for the Zelda close to the *Krüppel* core promoter. Zelda is an activator of CD1, and this site is likely to be bound *in vivo* since its predicted location coincides with a strong Zelda ChIP-seq peak ((Harrison et al., 2011) and K.B., unpublished data). We suspect that this may account for the weaker expression from their CD1-only construct compared to ours.

### CD2 represses CD1 through an unknown long-range mechanism

The CD2 enhancer represses CD1 activity in the posterior of the *Krüppel* pattern ((El-Sherif and Levine, 2016) and Figure 5A). We found that this repression does not depend on CD2 being next to the promoter or being positioned between CD1 and the promoter: CD2 represses CD1 just as effectively when the enhancer’s locations are switched (Figure 5B,C). The posterior border of *Krüppel* expression is set by Giant and Knirps, both of which are short-range repressors (Arnosti et al., 1996; Strunk et al., 2001). Short-range repressors act over distances of less than ∼100bp to quench the activity of nearby activators or inhibit other enhancers by directly repressing the promoter from within this range (Arnosti et al., 1996; Hewitt et al., 1999). Even in its endogenous position, CD2 is ∼200 bp away from the transcription start site, and when the positions of the enhancers are switched it is ∼2.4 kb away, meaning that short-range repressive action of Knirps or Giant cannot account for its inhibition of CD1. Repression between the *Krüppel* enhancers must therefore either be mediated by an unknown long-range repressor, or depend on previously undetected long-range repressive activity of Knirps and/or Giant.

### The effect of enhancer order suggests a role for topology in combinatorial control at the locus level

The constructs containing the shadow enhancer pair in their endogenous and switched locations are comprised of the same regulatory sequences, and yet they perform different computations to determine the overall gene expression output (Figure 5E). Topology affects gene expression at the genome scale, organized by DNA-binding looping factors whose position relative to one another is critical (Guo et al., 2015). Futhermore, gene expression is affected when topological interactions are forced through introduction of looping factors (Bartman et al., 2016; Deng et al., 2014). We hypothesize that looping factors may also influence gene expression at the level of single loci. In our constructs, these looping factors could be present either in the enhancers themselves or in the spacer sequences that replaced them in the single-enhancer constructs, though we deliberately designed the spacers to lack binding sites for many regulatory proteins (see Methods and Table S2). Indeed, predicted binding sites for multiple Drosophila looping proteins are present throughout our constructs (see Figure S2); investigating the role of these sites in locus-level computations is an exciting direction for future work.

### Center of pattern supports simple model of enhancer competition

Across the *Krüppel* pattern, but particularly at the center, pairs of enhancers tend to combine additively when their expression is low, and sub-additively when it is higher (Figure 3D, Figure 4). This is consistent with high expression levels being the result of increased enhancer-promoter contact, which would lead to competition between the enhancers if they can only activate the promoter one at a time (Bothma et al., 2015).

The levels of the repressors that set the boundaries of *Krüppel* expression are low in the center of the pattern, meaning that expression there is dictated primarily by the activators. An interesting feature of the *Krüppel* shadow enhancers is that they are controlled by different activators: CD1 is sensitive to Zelda and Bicoid (Hoch et al., 1991; Wunderlich et al., 2015), while CD2 responds to Stat92E and (perhaps indirectly) to Hunchback (Jaeger, 2010; Wunderlich et al., 2015). We had wondered whether CD1 and CD2’s use of distinct activators affects how their activities combine. If this were the case, we would expect the shadow enhancer pair to carry out a different computation to the two sets of duplicated enhancers. Instead, the shadow enhancers and duplications perform the same computation over a wide range of expression levels in the center of the pattern (Figure 3D), suggesting that the activator’s identities don’t matter in this case.

### Explaining differences in computation across the *Krüppel* expression pattern

The competition model as laid out by Bothma et al. (2015) cannot account for the combined output of two enhancers across the *Krüppel* expression domain as a whole. Specifically, having quantified expression across the entire *Krüppel* pattern, we observed significant differences in the computation performed at its anterior and posterior borders, both within and between constructs (Figure 4).

How might different combined ‘output’ levels be generated from the same summed ‘input’ level of expression when the two enhancers act independently? In the model described by Bothma et al., two enhancers can contact the same promoter one at a time and do not inhibit or assist one another’s activity. Achieving different computations under this model would require that the enhancers change the level of expression not just by changing the fraction of time that the enhancers engage with the promoter, but also by varying the rate of transcription initiation (Figure S3 and accompanying Supplemental Text). This runs counter to recent *Drosophila* studies that show enhancers altering the frequency of transcriptional bursts (changing the fraction of time that the promoter is active), rather than burst amplitude (i.e. the efficiency of transcription initiation during active periods) (Fukaya et al., 2016; Lammers et al., 2018; Zoller et al., 2017).

A simple explanation for the difference in computation at the anterior and posterior boundaries is that different TFs are present in each location.This explanation is supported by the behavior of the duplications because both enhancers respond to the same TFs but computations across the pattern still differ. Indeed, *Krüppel*’s anterior and posterior borders are determined by different combinations of transcriptional repressors. The anterior boundary is set by Giant and Hunchback and the posterior by Giant and Knirps (Harding and Levine, 1988; Jäckle et al., 1986; Jaeger, 2010; Kraut and Levine, 1991). Whether CD1 and CD2 respond differently to these repressors has not been directly tested. However, CD2’s expression pattern does not extend as far to the posterior as that driven CD1 and so it seems likely that CD2 is sensitive to Knirps (whose expression overlaps the posterior of *Krüppel*’s pattern) while CD1 is not. Transcription factors have different and context-dependent biochemical roles in transcription (Duarte et al., 2016; Fuda et al., 2009). In *Drosophila*, repressors can act on different steps in transcription when bound to shadow enhancers of the *sloppy-paired* gene (Hang and Gergen, 2017). A similar scenario may exist for *Krüppel*, with different combinations of repressors at the anterior and posterior of the pattern acting on distinct processes in transcription; the mechanism(s) by which each enhancer activates or represses transcription could in turn affect how the activities of two enhancers combine quantitatively.

### Outlook

Given the complexity of enhancer function over space and time, dynamic models of transcription are required to decipher the underlying molecular mechanisms. Models are most helpful when they can be confronted with relevant experimental data. While great strides have been made in measuring the dynamic production of mRNA using live imaging, measuring the underlying steps of transcription outside of bulk biochemical assays remains highly challenging. Single molecule approaches in bacteria have been very successful at linking models to mechanism through measurements of the underlying steps and their rates. Our results indicate that such measurements will also be transformative for animal systems, where the underlying biochemical rates likely influence not only how individual enhancers operate, but also how they collaborate with one another.

## ACKNOWLEDGEMENTS

This work was generously supported by the NSF Career Award –IOS-1452557 (AHD), NIH 4U01GM103804-04 (AHD), the Giovanni Armenise-Harvard Foundation (AHD) and a Harvard Graduate Society Merit Fellowship (CS). We would like to thank Hernan Garcia, who developed (and taught us to implement) both MS2 imaging for fly embryos and the pipeline we used to process the imaging data. We are grateful to Hernan for many helpful discussions. We thank Armando Reimer and Jacques Bothma for advice on imaging and image processing. We carried out all imaging at the Nikon Imaging Center at Harvard Medical School, where Anna Jost and Jennifer Waters provided invaluable guidance on microscopy. We thank members of the DePace lab for their input throughout this work.

## AUTHOR CONTRIBUTIONS

C.S. & A.H.D. conceived the study, designed the experiments and wrote the paper; C.S. conducted the experiments and analysed the data; K.B. helped make the transgenic fly lines; T.H. contributed to image processing and analysis.

## METHODS

### Cloning and transgenesis

We constructed a reporter gene in the pBOY vector backbone, into which we cloned the *Krüppel* (*Kr*) regulatory sequence out to ∼4.5 kb upstream of the transcription start site. The reporter consisted of the *D.melanogaster Krüppel* core promoter and its surrounding sequence from the 3’ end of the CD2 enhancer to the beginning of the second exon of *Krüppel*; a 1.5 kb cassette encoding 24 MS2 stem loops (from Addgene 31865, pCR4-24XMS2SL-stable plasmid); 3 kb of the *lacZ* gene; and the alpha-tubulin 3’ UTR. We commercially synthesised the 4.5 kb of *Krüppel* regulatory sequence that contains the proximal and distal blastoderm enhancers (using GenScript’s gene synthesis service) in order to insert unique restriction sites flanking each enhancer, enabling us to replace each sequence in a modular fashion. We then ligation-cloned this sequence into our pBOY construct, directly upstream of the *Krüppel* promoter. For subsequent manipulations of the two *Krüppel* enhancers, we used the sequence definitions from Perry et al. (2011); the sequences and genome coordinates for the *Krüppel* enhancers we used are listed in Table S1.

In order to maintain the endogenous spacing between enhancers and promoter in the single-enhancer constructs we replaced each with an equivalent length of non-regulatory sequence. We computationally designed these sequences be depleted of binding sites for transcription factors active in patterning the blastoderm embryo, sites for architectural binding proteins, and core promoter sequences (Supplementary Table 1). To do so we used the online binding site removal tool, SiteOut (Estrada et al., 2016), beginning with a randomly-generated sequence of the appropriate GC content for Drosophila intergenic DNA (40.6%). SiteOut locates binding sites on the basis of their Position Weight Matrices using PATSER (stormo.wustl.edu) and a p-value threshold that we set at 0.003. It then removes sites iteratively using a Monte Carlo algorithm while maintaining the GC content specified. We commercially synthesised a length of binding site-depleted sequence, and from it PCR-amplified unique sections of the appropriate lengths to replace the CD1 and CD2 enhancers (1160 bp and 1586 bp, respectively; Table S1). These we cloned into our reporter plasmid in place of the enhancers using isothermal assembly (Gibson et al., 2009), which leaves scarless junctions.

We sequence-verified the enhancers and promoter of all reporter constructs prior to injection, and checked the length of the MS2 cassette by restriction digest. The pBOY backbone contains an attB site for phiC31-mediated site-specific recombination (Fish et al., 2007) and a mini-white gene for transformant selection. For each construct, BestGene Inc. (Chino Hills, CA) injected midi-prepped DNA into 200 embryos of Bloomington Stock BL8622, which contains the attP2 landing site on chromosome 3L (Markstein et al., 2008). All constructs are integrated into this same attP2 landing site. After the constructs were successfully integrated into the fly genome, we prepared genomic DNA, PCR-amplified the transgene and repeated the sequencing and restriction digest verification of the reporters. Doing so revealed that a single construct, the CD1 duplication, may have four extra MS2 stem loops (giving it 56 rather than 48 copies of GFP if all stem loops are fully bound by MCP-GFP). This does not change the main conclusion that we draw from analysis of the CD1 duplication, which is that the computation performed by the two copies of CD1 is different in the anterior and the posterior of the expression pattern. Conservatively correcting for the possible extra loops simply shifts the data in Figure 4B down without changing their positions relative to one another (data not shown). Note that in the construct containing the shadow enhancers in their switched location (CD2*-CD1*), the proximal enhancer (which is endogenously located ∼200 bp upstream of the promoter) is at 2.4kb upstream, compared to 2.8 kb in the corresponding single-enhancer control.

### Live imaging: embryo preparation and data acquisition

Virgin females of the line yw; His2Av-mRFP1; MCP-NoNLS-eGFP (Garcia et al., 2013) were crossed to males bearing each of our transgenic reporter constructs. At two hours after egg deposition, embryos were dechorionated in freshly-made 50% bleach for 1 minute and then mounted in halocarbon 27 oil between a semi-permeable membrane (Biofoile, In Vitro Systems and Services) and a coverslip (No. 1.5, 18 × 18 mm).

Live imaging was carried out at 20 second intervals for the first 50 minutes of nuclear cycle 14 (nc14, beginning ∼2 hr post-egg deposition). Each time-lapse series was registered to its position on the anterior-posterior axis by cross-correlation with a 20x image of the full embryo. Our field of view was narrower than the expression pattern of some of our constructs; we therefore report mean fluorescence data from multiple embryos for each transgenic line, with an n of at least 3 for each anterior-posterior bin.

Live imaging was carried out using a Yokagawa CSU-22 spinning disk confocal with Borealis modification (Spectral Applied Research) with a Hamamatsu ORCA-R2 cooled CCD camera, mounted on an inverted Nikon Ti microscope. The MCP-eGFP and Histone-mRFP1 were imaged using a 488 nm solid state laser with 525/50 nm emission filter, and 561 nm solid state laser with 620/60 nm emission filter, respectively. Laser lines were selected using an AOTF, and a 405/488/568/647 multi-band pass dichromatic mirror (Chroma) was used. Time-lapsed images were acquired using a Nikon Plan Apo 60x 1.4 NA oil immersion objective. Low-magnification images used to register the movie on the anterior-posterior axis of the embryo were taken with a Nikon Plan Apo 20x 0.75 NA objective. Acquisition was controlled using MetaMorph software (Molecular Devices). To enable quantitative comparison of expression levels between embryos, we normalized illumination intensity at the start of each imaging session by measuring the power of 488 nm laser exiting the objective lens, using an EXFO X-Cite XR2100 power meter and adjusting the laser AOTF to reach a set target value. A 10 um z-stack of 21 images spaced 0.5 um apart was taken every 20 seconds using a Prior NanoScanZ piezo-electric focus stage insert, an exposure time of 80 ms, and 2×2 camera binning, resulting in a pixel size of 212 nm. Flatfield images were taken under identical imaging conditions to those described above, using a slide of concentrated fluorescien sodium salt solution (M.A. Model & J.L. Blank, 2007). Using the full CCD chip with 60x magnification did not cover the full extent of the *Krüppel* pattern; we report mean fluorescence data from multiple embryos for each transgenic line, with at least 3-fold coverage across the pattern from different embryos.

### Analysis of live imaging data

The imaging data were processed in Matlab (Mathworks) as described in Garcia et al 2013; the scripts are available from the Garcia lab at UC Berkeley. The histone-RFP image stacks were maximum-projected at each time point, and the nuclei segmented. MCP-GFP images were corrected for uneven laser illumination by subtracting the camera offset (a flat value of 200 added to every pixel) and then dividing by an offset-subtracted, normalised flatfield image. Spots of transcription were located in each z-slice of each time point using a difference of Gaussians filter and associated in z to a given site of transcription (or ‘particle’). Each particle was then associated with its closest respective nucleus; in cases where more than one particle is detected in the vicinity of a nucleus, the brightest particle alone was kept. A 2D Gaussian was fitted to the brightest z-spot in each particle to determine the offset, which was used as an estimate of the local background fluorescence. The fluorescence of a particle was calculated by integrating the fluorescence over a fixed area of 9 pixels diameter centered on the 2D gaussian in the image in the z-stack containing the highest intensity value pixels in this region, and subtracting the estimated local background. This background fluorescence estimate dominates the imaging error (Garcia et al., 2013). Spots for which there was no peak in fluorescence in the z-axis in a given time point (i.e. where part of the spot was cut off) were discarded.

A single 20x image of each embryo was generated by automatically stitching together two 512 × 675 pixel images (pixel size 600 nm) of the anterior and posterior of the embryo in Matlab (Figure S1A,B). The anterior, posterior, dorsal and ventral poles of the embryo were manually assigned and the full embryo image used to align the 60x imaging region by cross-correlation using the histone-RFP channel. Each particle of fluorescence was then assigned its anterior-posterior and dorso-ventral coordinates and parsed into anterior-posterior bins of 2.5% embryo length (Figure S1C).

The borders of the *Krüppel* expression pattern were calculated by fitting a Gaussian to the expression pattern (the mean fluorescence in ‘on’ nuclei) for a given construct at each time point (Figure S1E). The anterior (posterior) border was determined to be the position of the Gaussian’s peak minus (plus) the half width of the Gaussian at half max (HWHM).

The initial transcription rate was calculated for individual particle traces by finding the maximum derivative of the time series prior to the initial peak in its expression in nc14 (Figure S1D). The mean transcription rate for a construct was determined by averaging that for all the particles in a given anterior-posterior bin across all embryos of that construct.

